# Accelerated discovery and optimization of a potent and selective MMP12 inhibitor with the machine learning platform CAMOLE-1

**DOI:** 10.64898/2026.10.21.755545

**Authors:** Calico Life Sciences LLC, Jeffmin Lin, Matthew A. Bratkowski

**Affiliations:** Calico Life Sciences LLC

## Abstract

Machine learning techniques are being readily utilized to accelerate drug discovery pipelines. Peptide therapeutics are an emerging modality that offer the advantage of increased potency with reduced toxicity that are amenable to machine learning discovery and optimization. This work describes the **C**alico **A**ll-Atom **Mol**ecule **E**ngine (CAMOLE-1), a machine learning platform for de novo peptide inhibitor design. CAMOLE-1 incorporates an all-atom diffusion model that enables rapid non-canonical peptide discovery and optimization, and integrates physics-based computational filtering, and biochemical testing. We applied CAMOLE-1 towards the discovery of peptide inhibitors of matrix metalloproteinases (MMPs), a target class that has essentially evaded therapeutic development. Starting from canonical amino acid-based design, an MMP12 peptide inhibitor was identified and further optimized to improve potency. From these initial peptides, we used our all-atom diffusion model to determine a predicted structure, allowing us to design a potent and selective inhibitor over the course of about one year and 270 synthesized peptides. Predicted structures of both canonical and non-canonical peptide inhibitors showed high fidelity to our experimentally determined peptide-bound crystal structures. To our knowledge, this is the first described effort to use machine learning techniques to iteratively drive peptide inhibitor design beyond initial hit discovery into the development of selective, potent lead peptide inhibitors with improved pharmacokinetic properties. Our study illustrates the power of machine learning-driven design of peptide inhibitors, an exciting emerging therapeutic modality.

## Introduction

Small molecules and antibodies are the most common types of approved therapeutic modalities, but they are not ideally suited for all targets. Small molecules are routinely formulated to be orally bioavailable but typically require a well-defined binding pocket to be effective. Small molecules can also produce off-target effects by inhibiting proteins that are structurally similar to the target protein or inhibiting critical off-target proteins. While antibodies bind to a larger target surface area and are typically very selective, they are mainly restricted to extracellular targets and need to be administered by injection. Peptides are a therapeutic modality that bridges advantages between small molecules and antibodies. Peptides are more modular than small molecules and thus have the potential to be tunable for selectivity. Peptides bind to a larger protein surface and are well-suited for proteins that do not feature a traditional druggable pocket. While stability and permeability are challenges for developing peptide therapeutics, orally bioavailable peptides offer a distinct delivery advantage over injectable only antibodies. Exciting recent progress in oral therapeutic peptide development includes pill forms of the GLP-1 agonist semaglutide on the market^1–4^ , and very recent FDA approvals of the orally bioavailable IL23R inhibitor icotrokinra^5^ and PCSK9 inhibitor enlicitide^6,7^ .

Traditionally, peptide inhibitors are developed from hits found from peptide libraries utilizing phage^8^ or mRNA display^9^ campaigns. Machine learning methods can instead be applied to accelerate the development of peptide therapeutics. Recent advances in artificial intelligence have shown the potential for machine learning to drive rapid development of novel biomolecular binders to proteins of interest, with promising experimental characterization mostly on antibody, mini-protein, and macrocyclic peptide design^10–15^ . The use of current models in the drug design setting is focused on small molecules and antibodies. These models tend to be ill-suited for the development of highly potent peptide inhibitors, and residue-level coarse graining techniques may hinder generalization beyond the 20 canonical amino acids. Modeling approaches that incorporate atomistic modeling of non-canonical entities into peptide inhibitor design may help to close this gap. To our knowledge, however, a comprehensive effort to develop machine learning-designed peptides beyond zero-shot design into drug-like entities with promising pharmacokinetic behavior has not yet been described.

In this report, we describe CAMOLE-1 (**C**alico **A**ll-Atom **Mol**ecule **E**ngine), our machine learning platform for peptide therapeutic discovery and development. CAMOLE-1 is powered by a design loop consisting of an initial canonical amino acid design pipeline with scoring and identification of the most promising initial seed hits, non-canonical amino acid optimization over synthetically accessible chemical space, physics-based computational modeling and scoring, and peptide synthesis (Fig. 1). The platform features a transformer-based diffusion model for the structure prediction of peptides with non-canonical amino acids bound to targets of interest. This diffusion model eschews the coarse-graining of molecules into residue-level tokens and instead models both proteins and non-canonical amino acid containing peptides at an all-atom level. CAMOLE-1 integrates physics-based computational filtering to remove poor quality peptides at an early stage and incorporates experimental testing followed by improvement of drug-like properties. As a proof-of-concept study evaluating the effectiveness of our platform, we generated potential lead peptide inhibitors for matrix metalloproteinases (MMPs) that are involved in numerous disease states including rheumatoid arthritis, various cancers, oral diseases, cardiovascular diseases, and the lung diseases asthma, emphysema, and chronic obstructive pulmonary disorder, among other disorders^16,17^ .

**Figure 1:**
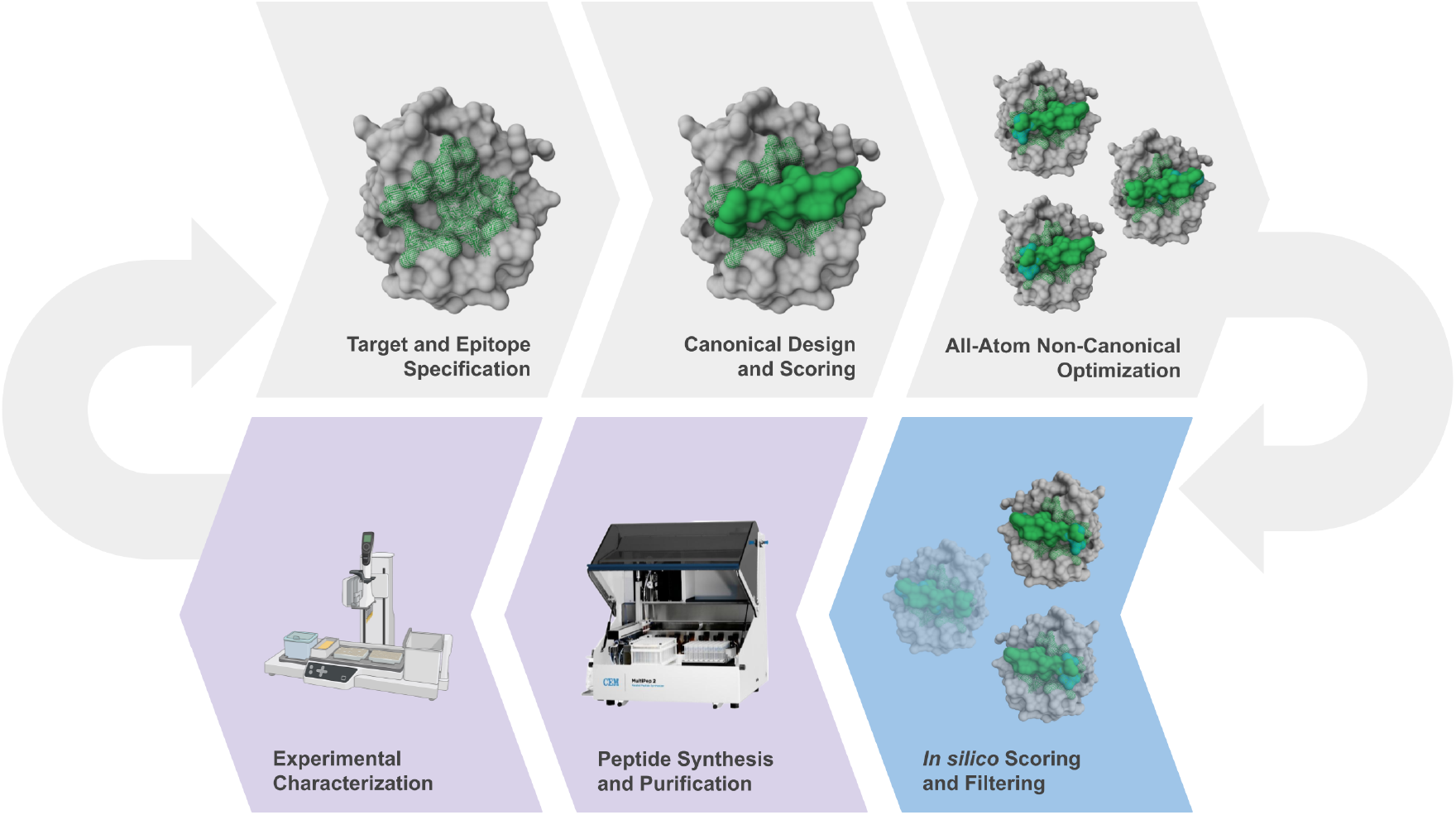
Overview of the platform and experimental characterization. Targets and hotspot residues are specified. A canonical amino acid-based machine learning design pipeline was used for initial hit generation. All-atom structural modeling enables rapid non-canonical optimization. Designed peptides are ranked and scored in silico with a combination of machine learning metrics (confidence, affinity) and physics-based simulation. Top candidates are synthesized and experimentally characterized, and data is fed back to the machine learning efforts.

MMPs are zinc bound proteases that cleave proteins in the extracellular matrix among other substrates including peptide hormones and chemokines^16^ . MMPs include the widely studied MMP1, 2, 3, 7, 8, 9, 10, 11, 12, 13 and 14 as well as other family members^16,17^ . Despite differences in individual MMPs, they all feature a similar, long catalytic groove with specific substrate sites classified using S and S’ (and corresponding substrate P and P’) protease nomenclature relative to the scissile bond^18^ . MMPs are not especially specific proteases because they cleave overlapping substrates and feature a high preference for leucine at the P1’ position^19^ . However, differences in the composition and depths of their subsites, especially the S1’ site, have been exploited for the design of more specific inhibitors^20,21^ . Lack of specificity of MMP inhibitors is an issue that still confounds small molecule inhibitor development^22,23^ . While there have been intensive efforts for over 30 years to develop selective MMP inhibitors, essentially none have passed clinical trials^21,24–26^ with the lone exception of Periostat, a low dose of the weak MMP inhibitor doxycycline that is used to treat the gum disease periodontitis^27,28^ . Peptide inhibitors serve as an attractive alternative modality compared to small molecules because they can be better modulated to the active subsites as suggested by several reported preclinical peptide-derived inhibitors for various MMPs^29–34^ .

CAMOLE-1 facilitated the development of a novel peptide inhibitor with an IC50 of 0.349 nM with promising pharmacokinetic properties that is at least 165-fold selective for MMP12 over other MMPs. This work demonstrates the use of CAMOLE-1 as a widely applicable approach to the rapid design and development of peptide inhibitors.

## Results

### *De novo* hit discovery and efficient optimization of MMP12 peptide inhibitors

MMPs are ideally suited for our machine-learning based peptide inhibitor design and experimental testing as a wealth of high-resolution crystal structures are available in the Protein Data Bank^35,36^ and a common fluorescence-based substrate probe facilitated straightforward activity assays^37^ . We focused on MMP2 and MMP12 as templates for model generation, and about 20 all-canonical amino acid peptides for each protein were synthesized. Due to the high similarity in active sites for all MMPs, we tested each peptide in activity assays for both proteins. Most peptides showed no activity inhibition, but we found one promising hit.

Peptide **1** emerged from this initial set of designs and was predicted to contain a short disulfide bonded, anti-parallel beta sheet with its N- and C-terminal residues close to the catalytic zinc atom (Fig. 2a). Peptide **1** inhibited 42% of MMP12 activity at the highest tested peptide concentration but did not inhibit MMP2 (Fig. 2b). Peptide **1** was selected for further optimization. We hypothesized that addition of the zinc chelating group hydroxamate would improve potency as this is a common feature of MMP inhibitors^38^ . To precisely incorporate this motif, we applied our physics-based computational pipeline to identify an optimal attachment site. Structural and energetic profiling revealed that truncating the C-terminal valine and substituting the adjacent threonine with threonine hydroxamate achieved ideal zinc coordination geometry. Crucially, the hydroxamate is predicted to engage the catalytic zinc while strictly maintaining the binding framework of the original machine learning-predicted peptide (Fig. 2c). We synthesized the hydroxamate-containing derivative of peptide **1** and refer to it as peptide **2**. Peptide **2** inhibited MMP12 completely with an IC50 of 154 nM while inhibiting MMP2 with an IC50 >100 µM (Fig. 2b and Table 1). Peptide 2 showed high selectivity for MMP12 over MMP1, 2, 8, 9, and 13, but low selectivity for MMP3, 7, and 10 (Table 1). Thus, peptide **2** proved to be a promising candidate for further potency improvement that was derived from a hit from our CAMOLE-1 platform.

**Table 1:** Peptide potency and selectivity data. Structures of macrocyclic hit peptide **1**, peptide **2**, and optimized peptide **13** with key amino acid changes in red and hydroxamate in blue (*top*), and IC50 potencies (*bottom*) in micromolar for peptide inhibitors against the catalytic domains of tested MMPs. Here and throughout the text, standard canonical amino acids are indicated by their one letter codes. “Thx” and “Ahx” represents threonine and alanine hydroxamate, respectively. The peptides are disulfide-bonded macrocycles. IC50s for curves that did not reasonably level off at 100% inhibition are labeled as >100 µM which is the highest peptide concentration used in the assay. “ND” indicates no data.

| Peptide | Sequence | MMP12 | MMP1 | MMP2 | MMP3 | MMP7 | MMP8 | MMP9 | MMP10 | MMP13 |
| --- | --- | --- | --- | --- | --- | --- | --- | --- | --- | --- |
| **1** | PVGTCGDDWCTV | >100 | ND | >100 | ND | ND | ND | ND | ND | ND |
| **2** | PVGTCGDDWC[Thx] | 0.154 | >100 | >100 | 0.000610 | 0.0125 | >100 | >100 | 0.0163 | >100 |
| **3** | AVGTCGDDWC[Thx] | 0.111 | ND | ND | ND | ND | ND | ND | ND | ND |
| **4** | PAGTCGDDWC[Thx] | 0.312 | ND | ND | ND | ND | ND | ND | ND | ND |
| **5** | PVATCGDDWC[Thx] | 0.0676 | ND | ND | ND | ND | ND | ND | ND | ND |
| **6** | PVGACGDDWC[Thx] | 0.192 | ND | ND | ND | ND | ND | ND | ND | ND |
| **7** | PVGTAGDDWC[Thx] | 24.7 | ND | ND | ND | ND | ND | ND | ND | ND |
| **8** | PVGTCADDWC[Thx] | 4.8 | ND | ND | ND | ND | ND | ND | ND | ND |
| **9** | PVGTCGADWC[Thx] | 0.245 | ND | ND | ND | ND | ND | ND | ND | ND |
| **10** | PVGTCGDAWC[Thx] | 0.463 | ND | ND | ND | ND | ND | ND | ND | ND |
| **11** | PVGTCGDDAC[Thx] | >100 | ND | ND | ND | ND | ND | ND | ND | ND |
| **12** | PVGTCGDDWC[Ahx] | 0.23 | ND | ND | ND | ND | ND | ND | ND | ND |
| **13** | PVGTCGPGWC[Thx] | 0.0099 | 6.72 | 0.823 | 0.0503 | 0.0929 | 1.09 | 2.21 | 0.154 | 4.63 |
| **14** | PVGTCGGGWC[Thx] | 0.234 | ND | ND | ND | ND | ND | ND | ND | ND |
| **15** | PVGTCGGPWC[Thx] | 9.5 | ND | ND | ND | ND | ND | ND | ND | ND |
| **16** | PVGTCGPPWC[Thx] | 21.44 | ND | ND | ND | ND | ND | ND | ND | ND |

**Figure 2:**
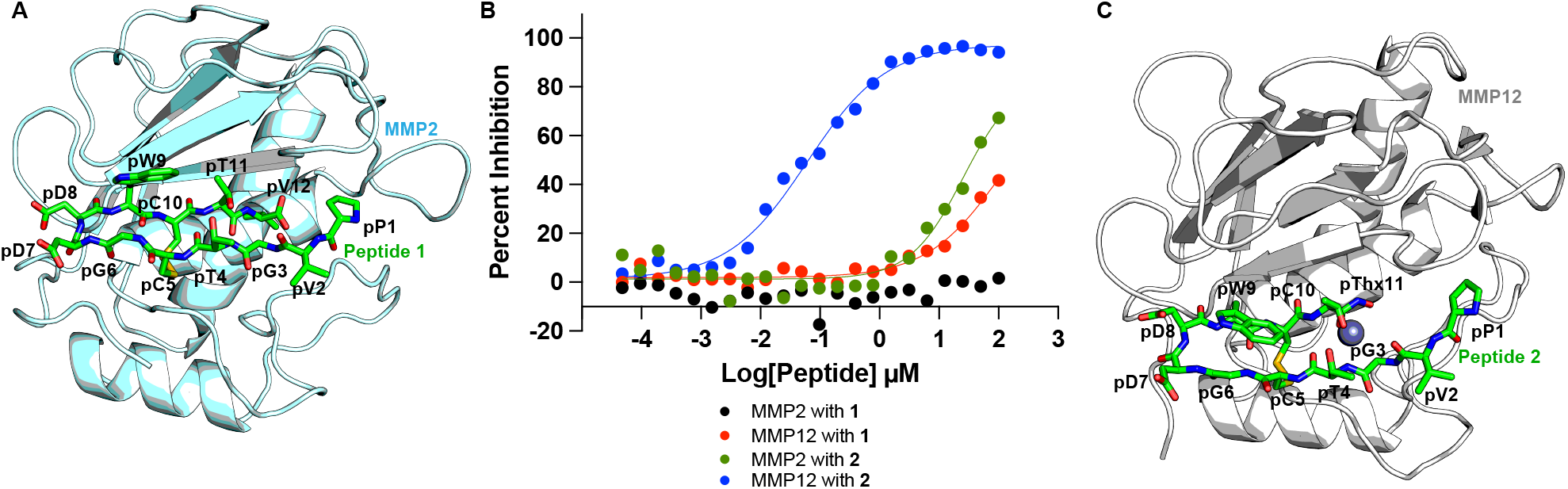
Machine learning-based peptide inhibitor hit finding and initial potency optimization. A) Machine learning model of MMP2 bound to peptide **1**. The published crystal structure of the catalytic domain of MMP2 with the fibronectin domains removed (PDB code 7xjo)^29^ was used as a template for machine learning. Here and throughout the paper, residues for the peptide are denoted “p” to differentiate them from protein residues. B) Fluorescence probe-based inhibition assay for peptide inhibitors against the catalytic domains of MMP2 and MMP12. C) MD-simulation derived structure that was used to model the interaction of MMP12 with peptide **2**.

### Potency optimization, crystal structure determination, and selectivity testing

Alanine scanning was conducted on peptide **2** to determine residues critical for potency (peptides **3** – **12**, Table 1). Replacement of most residues was tolerated except for pW9 (peptide **11**) that abolished potency, and pC5 (peptide **7**) and pG6 (peptide **8**) that significantly reduced it. Replacement of either of the two aspartates (peptides **9** and **10**) reduced potency by 1.6 – 3-fold. Analysis of the model (Fig. 2c) revealed that the aspartates are predicted to occupy a highly solvent-exposed segment forming a turn between two beta strands. To restrict the conformational ensemble, we hypothesized that replacing this segment with a canonical Pro-Gly (PG) sequence, a well-established motif for stabilizing beta-turns, would successfully rigidify the peptide framework. Computational modeling confirmed that the PG-substituted peptide retained the binding mode of the parent molecule (Fig. S1a) and MD simulations demonstrated that the PG substitution significantly reduced backbone flexibility, pre-organizing the peptide and minimizing the entropy penalty of binding (Fig. S1b). We synthesized this peptide (referred to as peptide **13**) and it had an improved potency of about 10 nM—about 15-fold more potent than peptide **2** (Table 1). In contrast, the peptide with replacement of glycine-glycine (peptide **14**) showed a similar potency to peptide **2**, and glycine-proline (peptide **15**) and proline-proline (peptide **16**) substitutions showed high micromolar potencies (Table 1).

In parallel with further machine learning-guided optimization (discussed below), we determined the crystal structure of **13** bound to the catalytic domain of MMP12 to 1.86 Å resolution (Fig. 3a and Table S1). As predicted, **13** forms a beta turn consisting of strand 1 (residues p1 – p7) and strand 2 (residues p9 – p11) with clear electron density observed for the peptide except N-terminal residues pP1 and pV2 (Fig. 3a). The beta turn structure of peptide **13** is formed by intramolecular hydrogen bonds between backbone atoms while the hydroxamate cradles zinc in a horseshoe-shaped trigonal-bipyramidal geometry (Fig. 3b) as observed previously^39^ . The hydroxyl of pThx11 hydrogen bonds to the catalytic residue E219 and the nitrogen hydrogen bonds to the carbonyl oxygen of A182. Strand 2 of **13** makes a continuous anti-parallel beta sheet with MMP12 residues 182-186 (“strand 3”) using backbone intermolecular hydrogen bonds. Notably, the side chain of H172 makes a π-π stacking interaction with pW9. Peptide **13** residue pG3 is bordered by P238 while the other strand 1 residues are outside of the catalytic groove and are partially solvent exposed, although some make packing interactions in the crystal structure (Fig. S2). Residues pThx11, pC10, and pW9 of strand 2 of peptide **13** occupy the S1, S2, and S3 pockets of MMP12 (Fig. 3c). While 70 crystal structures of MMP12 in complex with inhibitors have been published^40^ , peptide **13** is a novel inhibitor because it only binds in the S subsites while other inhibitors predominantly bind the S’ subsites and typically feature a functional group that binds in S1’ site tunnel (Fig. S3). Overall, the crystal structure confirmed the ability of our platform to produce de novo hits that can be optimized into improved inhibitors.

**Figure 3:**
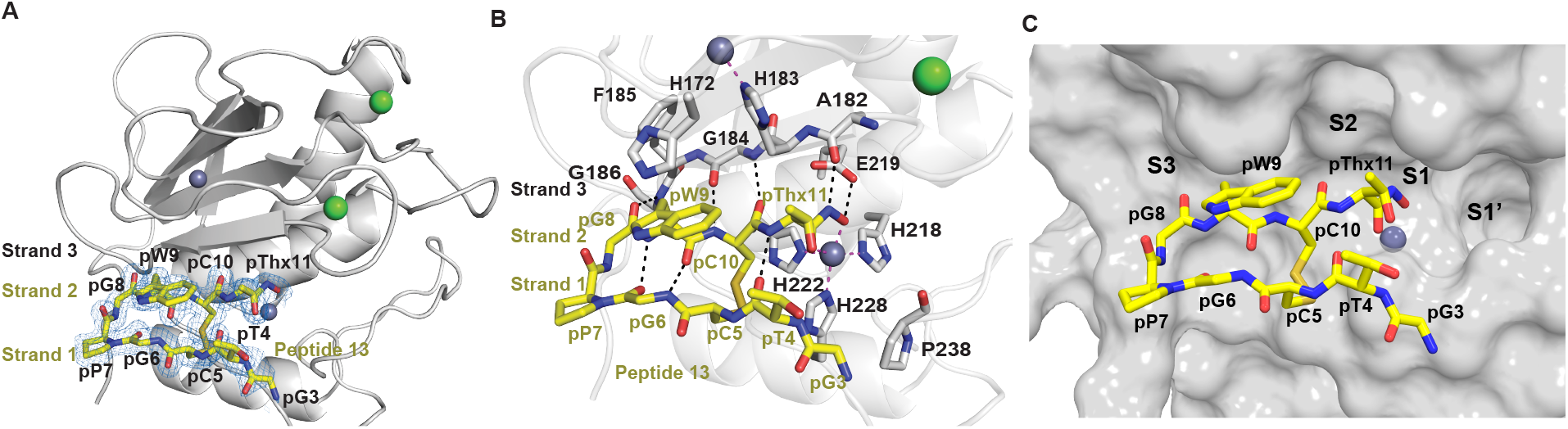
Crystal structure of MMP12 bound to peptide 13. A) Crystal structure of the catalytic domain of MMP12 (residues 106-263) bound to peptide **13** with 2FoFc electron density contoured at 1σ (*blue mesh*) overlaid on the peptide (*yellow sticks*). Here and throughout the report, zinc and calcium ions are shown as gray and green spheres, respectively. Hydrogen bonds are shown as black, dashed lines. B) Detailed interactions of **13** with MMP12. Metal coordination interactions are shown as magenta, dashed lines. The two strands of **13** make extensive backbone hydrogen bonds with each other and to “strand 3” of MMP12 to form a triple beta sheet conformation. C) Surface depiction of MMP12 with S3, S2, S1, and S1’ subsites labeled and peptide **13** in sticks.

We next tested the selectivity for peptide **13** against a panel of MMPs. Peptide **13** showed an improvement in selectivity for MMP3, 7, and 10 but a decrease in selectivity for MMP1, 2, 8, 9, and 13 compared to peptide **2** (Table 1). Selectivity of peptide **13** for MMP12 is likely partially dictated by residue H172 as it interacts with pW9 (Fig. 3b) but is not conserved in other tested MMPs (Fig. S4). Replacement of this residue by serine in MMP1 and 8 (Fig. S5a), and by tyrosine in MMP3, 7, and 10 (Fig. S5b and Fig. S5c) may partially explain the respective high and low selectivity for these groups (Table 1). We were not able to obtain a crystal structure of MMP12 containing “active site edge” residues 100 – 105 bound to peptide **13**, but alignment with a longer MMP12 structure^41^ indicates that non-conserved residue M103 could contribute to peptide binding differences in this region and hypothetically explain selectivity (Fig. S5a, Fig. S5b, and Fig. S5c). However, traditional structure-based SAR for potency and selectivity optimization would be a complex process and it is possible that any gains in selectivity for one MMP could easily be lost in another as was already observed upon optimization from peptide **2** to **13** (Table 1). To avoid such pitfalls, we further leveraged CAMOLE-1 for more efficient optimization using all-atom modeling with non-canonical amino acids.

### Rapid non-canonical optimization with an all-atom diffusion model

Accurate modeling of biomolecular interactions plays a critical role in de novo drug design. Recent work has shown that incorporating a mix of residue-level features with atom-and-bond level features can enable folding models to predict the structures of complexes beyond proteins, including modified canonical residues, lipids, antibodies, DNA/RNA, and small molecules^42–47^ . Graph neural networks which model entirely at the atomistic level further endeavor to enable modeling of novel chemical material and capture conformational diversity^48,49^ . The improved capabilities of these models to capture more general atomistic interactions promises to empower machine learning-guided drug design pipelines to generate non-canonical amino acid and heteroatom-containing peptides that can be developed into novel therapeutics. To enable our platform to generalize beyond protein-small molecule interactions into peptide-scale non-canonical interactions, CAMOLE-1 incorporates an all-atom graph neural network diffusion model which operates directly on atom and bond features (Fig. 4a). We encode the structure of our candidate molecules as well as the entire target protein in a consistent way, with unified node and edge feature preparation. Our model utilizes a general form of node and edge attention and triangle multiplication, first introduced by Jumper et al.^50^ We removed triangle attention in favor of scaling the rest of the architecture and included bi-directional node-edge feature mixing. The reverse diffusion process directly operates on 3D coordinates of every atom. Compared to co-folding models with specialized input tracks for canonical amino acids^42–47^ , by avoiding coarse-graining of molecules at a residue level, our all-atom featurization allows us to generalize to peptide-scale highly non-canonical chemical interactions. Using our all-atom modelling capabilities, we predicted the structure of MMP12 bound to peptide **13** and used it immediately for further optimization prior to completing our crystal structure. Notably, our predicted all-atom-based structure is in very good agreement with the crystal structure (RMSD = 0.664 Å for the peptide, and 0.343 Å for the whole structure), indicating a high level of fidelity for our system (Fig. 4b).

**Figure 4:**
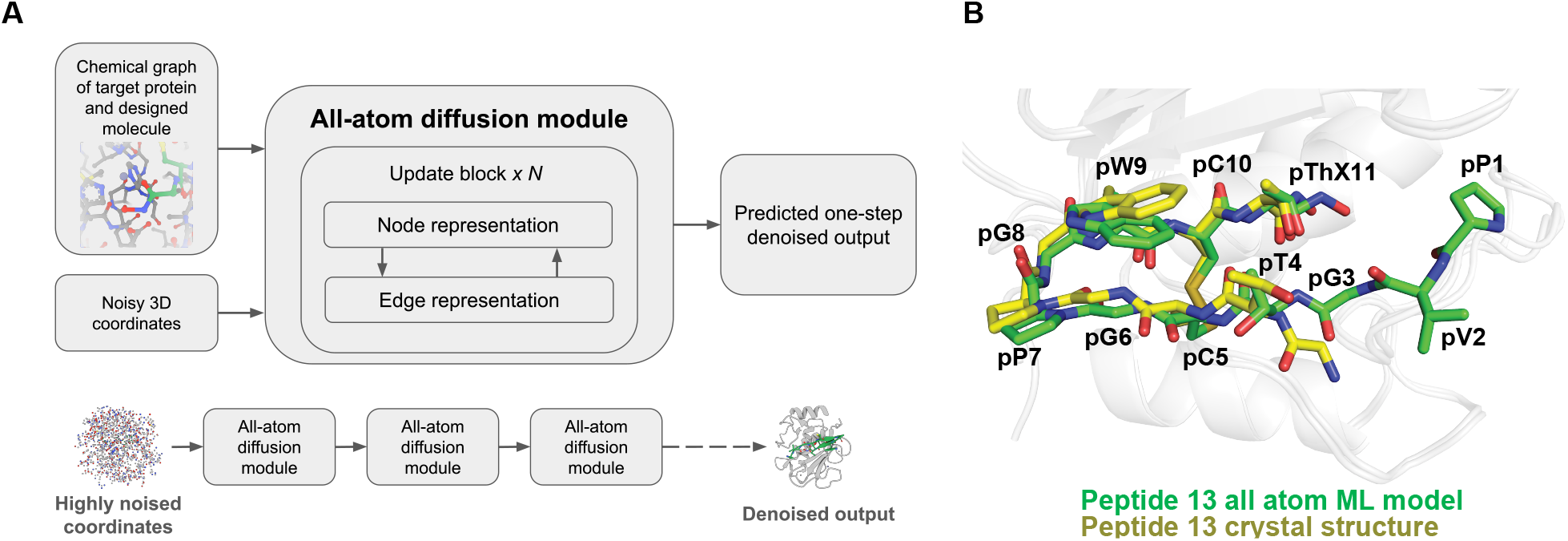
All-atom diffusion modeling integration into the pipeline. A) The diffusion module takes as input the atom-and-bond chemical graph of the target protein and binder, and after update blocks which bidirectionally mix node and edge features, the predicted noise is used in a reverse diffusion process to generate predicted 3D coordinates for all atoms. B) Alignment of the crystal structure of MMP12 bound to peptide **13** (*yellow sticks*) with the model prediction (*green sticks*).

Using our predicted structure directly for optimization, we generated thousands of candidates optimized from peptide **13** and selected 715 peptides for further scoring and computational filtering. Residues pC5, pG6, pW9, pC10, and pThx11 were retained in the same relative positions while the other residues were allowed to vary. Varied residues sampled canonical L-amino acids, D-amino acids, and other non-canonical amino acids. Candidate peptides were filtered for chemical feasibility using our physics-based computational pipeline. Initial structures were screened to eliminate unphysical geometries and subjected to orthogonal molecular docking to validate predicted binding poses. Candidates were then prioritized for synthesis based on ensemble thermodynamic scoring, clustering based on robustness and closeness to predicted poses, peptide conformational strain analysis, and physicochemical profiling.

We selected 94 peptides for synthesis and tested them for potency. Peptides displaying IC50 values of around 10 nM (Table S2) or better were analyzed for further work. Potent designs highly favored replacement of the first four residues (PVGT) from peptide **13** with XX G/dS A/G/dS, where X is any residue. Notably, these changes are not obvious from analysis of our peptide **13** bound crystal structure as residues pP1 and pV2 are unresolved (Fig. 3a). Potent peptides featured various residues in position 8 that included G, dS, dI, and dH. Residue dP was heavily favored in position 8 (77% of peptides) with the prominent sequence XX GACGP dP WC Thx (36% of peptides). Potencies for this series were improved compared to peptide **13** and ranged between about 3 – 8 nM (Table 2). Peptides **33** (SVGACGP dP WC Thx) and **38** (SIGACGP dP WC Thx) had the best potencies in this series with IC50s of 1.19 and 2.52 nM. Peptides **20, 25, 27, 32, 39**, and **52** that contained dP in position 8 also had IC50s of less than 3 nM. Other high potency peptides were **51** that featured dS in position 8, and **30** and **50** that had G in this position.

**Table 2:** Potency and selectivity data for all-atom model optimized peptides. Structure of peptide **51** with D-serine in green and hydroxamate in blue (*top*), and comparison of IC50 potencies in micromolar between catalytic domains of tested MMP proteins in the probe-based assay for peptides derived from all atom-based screening. Here “d” indicates that a D-amino acid was used in the peptide.

| Peptide | Sequence | MMP12 | MMP1 | MMP2 | MMP3 | MMP7 | MMP8 | MMP9 | MMP10 | MMP13 |
| --- | --- | --- | --- | --- | --- | --- | --- | --- | --- | --- |
| **20** | ATGACGS[dP]WC[Thx] | 0.00294 | 81.8 | 24.3 | 2.53 | 2.47 | >100 | >100 | 1.23 | >100 |
| **25** | [dL]PGGCG[NMeA][dP]WC[Thx] | 0.00229 | 8.18 | 1.04 | 0.697 | 0.274 | 16.6 | 8.92 | 0.219 | 60.9 |
| **27** | SP[dS]ACGP[dP]WC[Thx] | 0.00146 | >100 | >100 | 2.57 | 1.84 | >100 | >100 | 1.27 | >100 |
| **30** | TAG[dS]CGPGWC[Thx] | 0.00182 | 4.98 | 2.04 | 0.104 | 0.226 | 2.77 | >100 | 0.0791 | 32.2 |
| **32** | DVGGCGP[dP]WC[Thx] | 0.00101 | >100 | 48.5 | 0.336 | 1.31 | 41.6 | >100 | 1.03 | 40.7 |
| **33** | SVGACGP[dP]WC[Thx] | 0.00119 | >100 | >100 | 0.882 | 1.15 | >100 | >100 | 1.19 | 12.7 |
| **38** | SIGACGP[dP]WC[Thx] | 0.00252 | >100 | 61.4 | 0.601 | 0.666 | >100 | >100 | 0.619 | 12.8 |
| **39** | [Cval][Hyp]GGCGP[dP]WC[Thx] | 0.00284 | 97.5 | 12.4 | 0.615 | 0.889 | >100 | 50.8 | 0.289 | 20 |
| **50** | A[Hyp]GGCG[NMeT]GWC[Thx] | 0.00282 | 1.1 | 2.11 | 0.584 | 0.666 | 1.14 | 11.7 | 0.470 | 41.1 |
| **51** | [dS]PGACGP[dS]WC[Thx] | 0.00221 | >100 | >100 | 0.927 | 1.58 | 33.4 | >100 | 2.09 | >100 |
| **52** | APGGCG[NMeA][dP]WC[Thx] | 0.00292 | 10.8 | 1.84 | 0.566 | 0.584 | 47.6 | >100 | 0.437 | 26.8 |

The aforementioned peptides were shortlisted for further selectivity testing. Many peptides showed IC50s of >100 µM for MMP1, 2, 8, and 9 but much less selectivity for MMP3, 7, and 10 (Table 2). Peptides **25, 50**, and **52**, which all contained N-methylated amino acids, as well as peptide **30** inhibited MMP1 with IC50s of about 10 µM or less and MMP2 with IC50s of about 2 µM and were thus excluded from further work. Peptides **20, 27**, and **51** displayed good selectivity profiles and were selected for further optimization. Peptides **32, 33**, and **38** had very similar sequences and potencies so only **33** was moved forward for additional studies because it had a slightly better selectivity profile.

We wanted to perform a sanity check of our all-atom model with the machine learning-optimized peptides, so we determined a crystal structure of MMP12 bound to 51 to 2.59 Å resolution that featured clear density for the peptide (Fig. 5a and Table S1). The majority of peptide **51** assumes a similar orientation compared to peptide **13** (Fig. 5b) and the same molecular interactions are observed (Fig. S6). While the N-terminus of peptide **51** is slightly more resolved than peptide **13**, this may be due to crystal packing artifacts (Fig. S7), and no additional interactions were observed. The prediction from our all-atom diffusion model was in good agreement with the crystal structure (RMSD = 2.512 Åfor peptide and RMSD = 0.496 Å for the whole structure) except for the N-terminus (Fig. 5c), highlighting the validity of using the all-atom model for inhibitor optimization. While the structures of both peptides bound to MMP12 are similar, peptide **51** is about 4.5-fold more potent than peptide **13** and more selective (Table 2) and this is not obvious from analyzing the crystal structures alone. Overall, our all-atom platform allows for a heightened degree of optimization.

**Figure 5:**
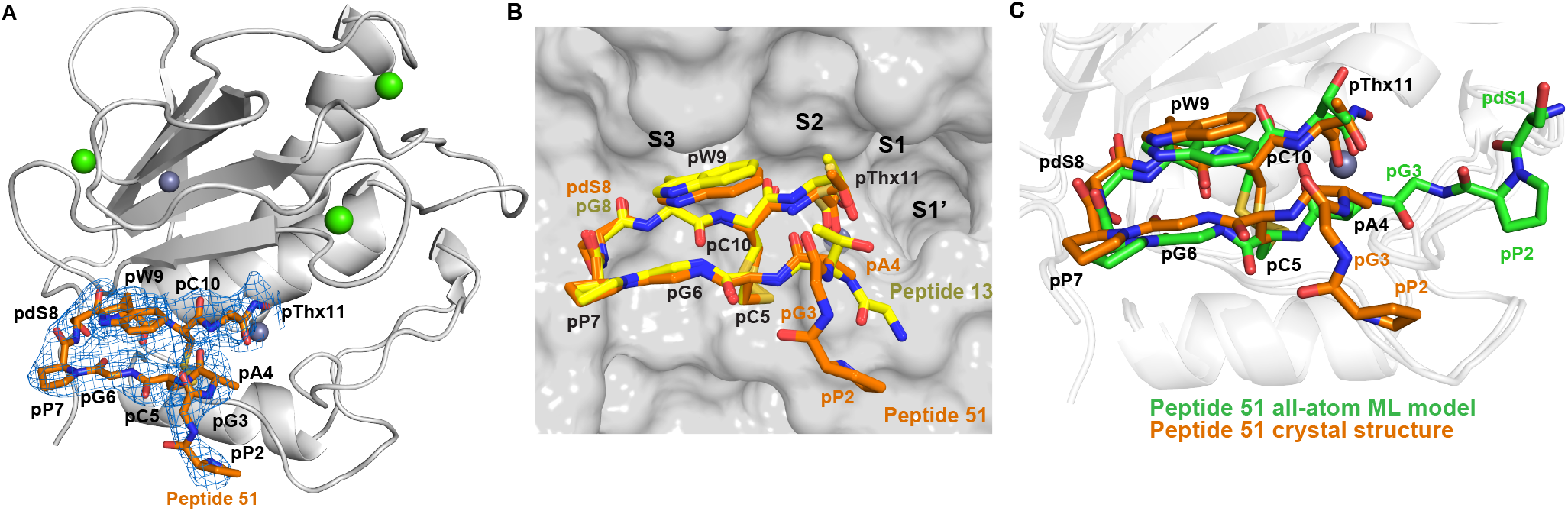
Validation of the peptide 51 bound model. A) Crystal structure of MMP12 bound to peptide **51** with 2FoFc electron density contoured to 1σ (*blue mesh*) overlaid on the peptide (*orange sticks*). B) Alignment of the crystal structures of MMP12 bound to peptide **13** (*yellow*) and peptide **51** (*orange*). C) Alignment of the MMP12 bound peptide **51** crystal structure (*orange*) with the all-atom machine learning model (*green*).

### Stability enhancement for improved drug-like properties

Peptides often need to be modified to be stable in vivo for use as therapeutics as they would otherwise be degraded by endogenous proteases. Acetylating the N-terminus of peptides prevents them from degradation by aminopeptidases. We therefore tested acetylated forms of peptides **20, 27, 33**, and **51**. Unexpectedly, acetylation decreased potency by 7-fold for peptide **56** (acetylated peptide **20**), 13-fold for peptide **57** (acetylated peptide **27**), and about 3-fold for peptide **58** (acetylated peptide **33**) compared to the parent molecules, but peptide **59** (acetylated peptide **51**) had no potency decrease compared to its parent peptide; all acetylated peptides retained a similar selectivity profile compared to their parents (Table 3).

**Table 3:**
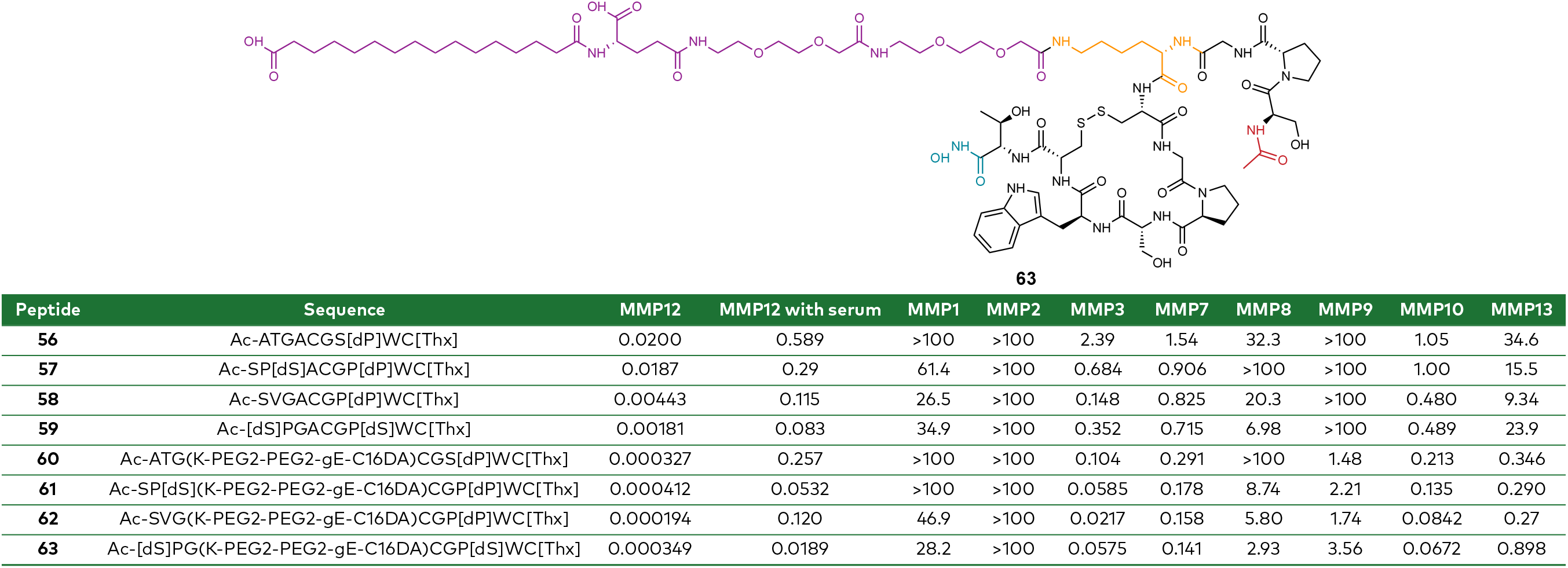
Potency, serum shift, and selectivity data for acetylated and lipidated peptides. Structure of peptide **63** (*top*) with acetyl cap (*red*), added lysine (*orange*), PEG2-PEG2-gE0C16DA (*purple*), and hydroxamate (*blue*) color-coded, and IC50 potencies in micromolar (*bottom*).

The addition of a lipid group was previously shown to extend the serum half-life of GLP-1 agonists^51–55^ so we applied this modification to our peptides. Based on our machine learning prediction of peptide **51**, we chose the solvent exposed third residue in the sequence (dS for peptide **57** and G for the other parent peptides) as an appropriate site for lipidation. Following previously reported lipidation strategies^56^ , lysine appended with a PEG linker and diacidic fatty acid (PEG2-PEG2-gE-C16DA) was installed at position 3 to produce peptides **60** (lipidated peptide **56**), **61** (lipidated peptide **57**), **62** (lipidated peptide **58**), and **63** (lipidated peptide **59**). Unexpectedly, lipidation increased the potency for all peptides to between 0.194 – 0.412 nM (Table 3). Since introduction of hydrophobic groups can increase non-specific interactions with proteins, it was important to test selectivity. All lipidated peptides showed high selectivity against MMP1 and 2, but showed increased inhibition of MMP3, 7, 9, 10, and 13 compared to their parents (Table 3). However, considering the increased potency for MMP12, the fold selectivity was still reasonable.

Addition of lipids is a double-edged sword in that it can increase serum half-life but also can reduce potency due to sequestration by human serum albumin. We therefore tested the lipidated peptides in a serum-shift assay. As expected, all peptides showed a reduction in potency in the presence of serum (Table 3). Peptide **63** showed the least potency decrease in serum (about 50-fold reduction), so it was selected for pharmacokinetic profiling. Compared to the non-lipidated peptide **59**, peptide **63** showed significantly increased stability (Fig. 6) as well as about a 2-fold improved in vitro plasma half-life, a 4-fold improved in vivo half-life, and about a 50-fold reduced clearance (Table 4). Thus, peptide **63** is a promising lead peptide for future therapeutic development.

**Table 4:**
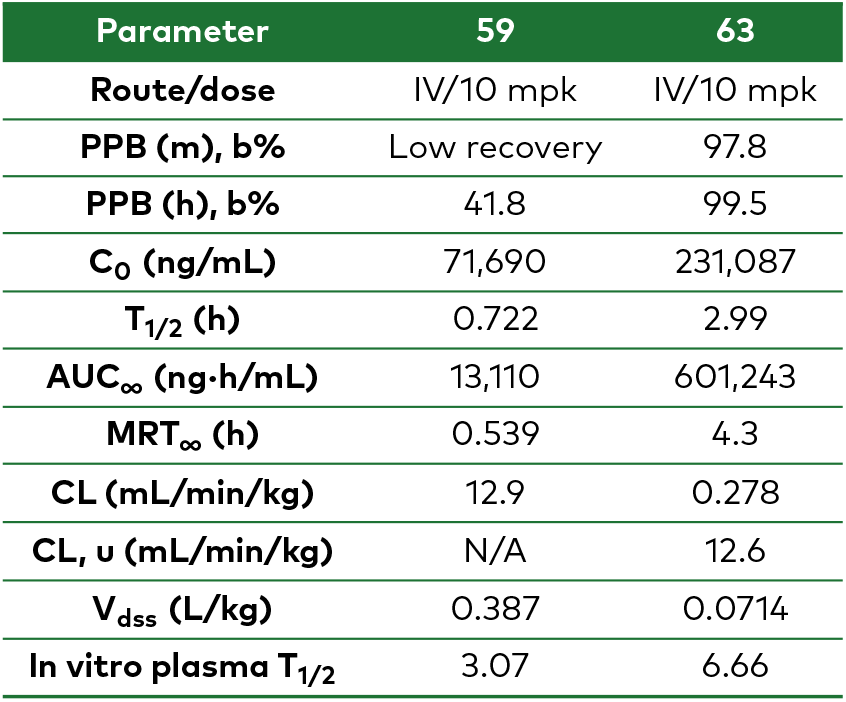
PK data for non-lipidated peptide 59 and lipidated peptide 63.

| Parameter | 59 | 63 |
| --- | --- | --- |
| Route/dose | IV/10 mpk | IV/10 mpk |
| PPB (m), b% | Low recovery | 97.8 |
| PPB (h), b% | 41.8 | 99.5 |
| C<sub>0</sub> (ng/mL) | 71,690 | 231,087 |
| T<sub>1/2</sub> (h) | 0.722 | 2.99 |
| AUC<sub>∞</sub> (ng·h/mL) | 13,110 | 601,243 |
| MRT<sub>∞</sub> (h) | 0.539 | 4.3 |
| CL (mL/min/kg) | 12.9 | 0.278 |
| CL<sub>u</sub> (mL/min/kg) | N/A | 12.6 |
| V<sub>dss</sub> (L/kg) | 0.387 | 0.0714 |
| In vitro plasma T<sub>1/2</sub> | 3.07 | 6.66 |

**Figure 6:**
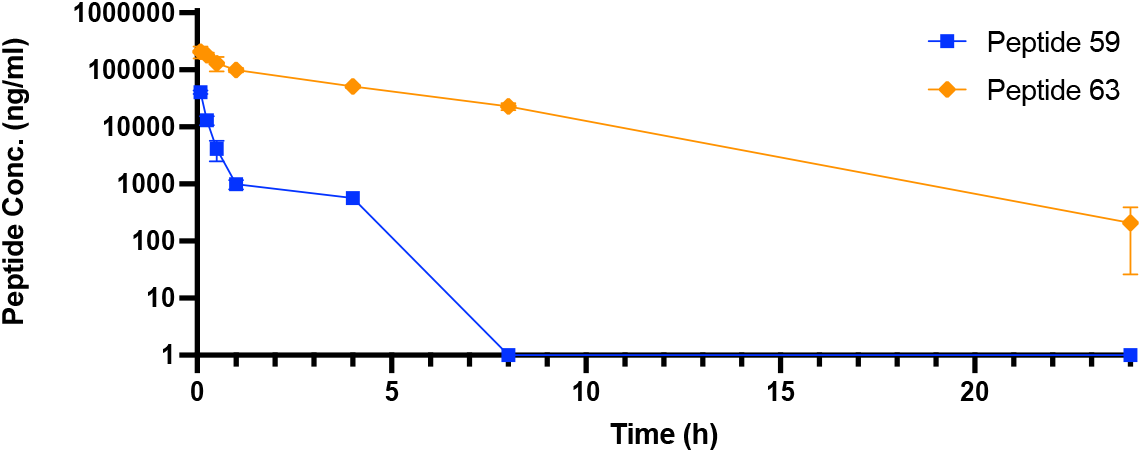
Stability test for peptides. A 24-hour pharmacokinetic profile for peptides **59** and **63** by 10 mg/kg intravenous (IV) dosing in mice. The concentration of peptide remaining is plotted at measured time points.

## Discussion

CAMOLE-1 is a machine-learning driven peptide inhibitor design platform that integrates both computational and experimental techniques to rapidly discover and develop potent and selective inhibitors. Our de novo canonical design campaign generated a hit with weak potency and low selectivity (peptide **1**) that was quickly optimized using rational design for potency for MMP12 (peptide **13**). Our all-atom-based diffusion model empowered us to generalize beyond canonical amino acid design. Accurate all-atom structure prediction unlocked rapid hit-to-lead optimization that was not obvious from expert crystal structure analysis. Additional optimization for stability resulted in peptide inhibitor **63** with sub-nanomolar potency and promising pharmacokinetic properties. Our platform allowed us to rapidly progress from a low potency hit to a peptide inhibitor with favorable properties for further therapeutic development over the course of about one year and 270 synthesized peptides. To our knowledge, this is the first report where machine learning methods have been applied to the discovery of a potent and selective MMP12 peptide inhibitor.

MMP12 is a target of great therapeutic interest as its overexpression is associated with the lung diseases emphysema, asthma, COPD, and allergic airway inflammation^57–59^ ; the cardiovascular diseases ischemic stroke and coronary artery disease^60,61^ ; and several cancers^62–65^ . A multitude of MMP12 inhibitors have been developed, including peptide-like molecules^40^ . Notably, the small molecule inhibitor Aderamastat has shown favorable results in an allergic asthma model^66^ and has advanced to clinical trials. Our machine learning-derived peptide inhibitors, however, are unique because they achieve selectivity by binding exclusively in the more conserved S1 – S3 pockets and avoid the S1’ pocket that is typically considered necessary for selectivity.

The use of machine learning drug design tools was a distinct advantage that propelled our pipeline at a quick pace that could not have been achieved by other methods alone. While optimization from peptide **13** to **51** was driven solely by computational techniques, notably the predicted models are in very good agreement with the crystal structures. This is particularly an advantage for peptide therapeutic discovery as the high-throughput determination of protein-peptide complexes can be challenging. Importantly, machine learning drove peptide potency and selectivity despite a similar overall protein-bound conformation and essentially identical interactions revealed from the crystal structures.

Although we used CAMOLE-1 to develop a peptide inhibitor against MMP12, it can be applied to other therapeutic targets to quickly drive discovery efforts. CAMOLE-1 offers significant promise for targets that have been considered “undruggable” by traditional small molecule inhibitors. Moreover, progress in peptide formulation^67^ suggests that our platform will facilitate discovery of peptides that can be developed into orally bioavailable therapeutics that offer a significant delivery advantage over injectable biologics-based therapeutics. Our platform demonstrates the power of deeply integrating machine learning with experimental characterization and principles of medicinal chemistry structure activity relationship. It is a streamlined, iterative process, incorporating data from many modalities—including structure, affinity, activity, and molecular properties—into machine learning models that will unlock rapid developable molecule generation and represents the next frontier in rapid therapeutic design.

### Experimental Methods

#### Peptide Synthesis

Linear peptides were synthesized using standard Merrifield solid phase synthesis techniques on a Multipep 2 (CEM) (peptides **17-55**) or by manual synthesis (peptides **1-16**; **56-63**) using standard coupling reagents O-Benzotriazole-N,N,N’,N’-tetramethyluroniumhexafluorophosphate (HBTU, 4.9 equiv.), N,N-diisopropylethylamine (DIPEA, 10 equiv.), and protected amino acid (5 equiv.). Fmochydroxylamine-2-chlorotrityl resin (0.64 mmol/g, sc-479116, Santa Cruz Biotechnology) was used to generate hydroxamate C-termini while pre-loaded Wang Resin with N-α-Fmoc protected amino acids was used for peptides with C-terminal acids. Global deprotection and cleavage from resin was performed by addition of 95% TFA/2.5% TIPS/2.5% H2O, followed by ether precipitation or TFA evaporation with heated nitrogen.

Oxidative cyclization of the dried linear peptides was performed by dissolving the peptides in 50% acetonitrile in water at approx. 5 mM, then adding a solution of 50 mM iodine dropwise with mixing until a yellow color persists. Excess iodine was quenched with a dropwise addition of 1 M ascorbic acid in 50% acetonitrile/water until the solution became colorless. Lyophilization of the crude cyclic peptide was followed by resuspension in 20% acetonitrile in water with 0.1% TFA. The crude solution was heated, mixed, and filtered before purification by reverse phase C18 HPLC. Purified fractions were pooled and lyophilized to give a white powder. Final products were analyzed by LC-MS to confirm purity by LC (absorbance at 214 nm) and identity by mass spectrum. A threshold of 70% purity by LC was set for all compounds from Multipep synthesis and 90% for manual synthesis.

#### In vitro MMP inhibition assay

IC50 measurements were determined using a fluorogenic pan-MMP quench probe with a KPLGL sequence (R&D Systems #ES010) and commercial catalytic domain MMP proteins from Enzo Life Sciences: MMP1 (BML-SE180), MMP2 (BML-SE237), MMP3 (BML-SE109), MMP7 (BML-SE181), MMP8 (BML-SE255), MMP9 (BML-SE360), MMP10 (BML-SE329), MMP12 (BML-SE138), and MMP13 (BML-SE246). MMPs were run at a final concentration of 3 nM (or at 1.5 nM for MMP13) with a final substrate concentration of 940 nM in a buffer containing 50 mM HEPES pH 7.5, 10 mM CaCl_2_ , and 0.05% Brij-35. The final DMSO concentration for each reaction was normalized to 1%. Peptides were tested at a serial titration range from 100 µM to 97.7 nM initially. Potent compounds that measured outside the dynamic range of the initial titration were run at a titration range of either 10 µM to 9.77 nM or 1 µM to 0.977 nM, depending on the peptide potency. Peptides were preincubated with the MMP for 15 minutes before starting the reaction with substrate. Fluorescence intensity (λex: 320 nm, λem: 405 nm) was measured on a CLARIOstar Plus plate reader after a 30-minute reaction time, ensuring the reaction was still in the linear range. For each MMP, the reaction rate of the DMSO control was defined as 0% inhibition and the reaction rate without protein was defined as 100% inhibition. IC50 values were then determined by nonlinear regression of inhibition versus inhibitor concentration using a four-parameter logistic model using CDD Vault (Collaborative Drug Discovery, www.collaborativedrug.com) or Graph Pad Prism Software (www.graphpad.com). IC50 values were judged as >100 µM if curves did not reasonably level off at the highest tested peptide concentration of 100 µM.

#### In vitro MMP inhibition serum shift assay

Serum shift assessment on MMP activity was measured in a similar set up to the in vitro inhibition assays described above with the replacement of 10% of the buffer with 10% human serum (Fisher Scientific, Cat#BP2657100) during the peptide and MMP incubation step, and a far-red probe was used as the pan-MMP substrate (Enzo Life Sciences Cat #: BML-P277). All other assay conditions and analysis were maintained.

#### Pharmacokinetic experiments

Pharmacokinetic profiling of peptides was conducted by WuXi Apptec. For stability testing shown in Fig. 6, peptides were formulated at 2 mg/mL in saline and dosed intravenously at 10 mg/kg to separate groups of three mice. Plasma concentrations for each peptide were measured at 9 timepoints out to 72 hours total.

#### Protein Expression and Purification

To make the construct for the version of MMP12 used for crystallography studies, the coding sequence for the catalytic domain of human MMP12 (Uniprot ID P39900, residues G106-G263) was codon optimized for expression in *E. coli* and cloned into a pET28a vector preceded by an initial methionine (Genscript). Protein expression and purification was completed by Biortus. The protein was expressed in BL21(DE3) cells under kanamycin selection at 37 ºC and induced with 1 mM IPTG for 4 hours at the same temperature. A volume of 2 L of culture was harvested by centrifugation to produce a 6.8 g pellet.

A denaturing purification followed by refolding was used to purify MMP12. The cell pellet was resuspended in 40 mL of lysis buffer consisting of 20 mM Tris-HCl pH 8.5, 1 M NaCl, 5 mM beta-mercaptoethanol, and 10 mM EDTA. The resuspension was then lysed with a high-pressure microfluidics homogenizer at 750 Bar for 4 cycles. Lysed cells were then centrifuged at 16,000 rpm for 30 minutes at 4 ºC and the inclusion bodies were pelleted and collected. The supernatant was discarded and the inclusion bodies were washed once with Wash Buffer A (20 mM Tris-HCl pH 8.5, 5 mM beta-mercaptoethanol, 10 mM EDTA, and 0.5% Triton X-100) followed by centrifugation at 16,000 rpm for 10 minutes at 4 ºC. The inclusion bodies were then washed twice with Wash Buffer B (20 mM Tris-HCl pH 8.5, 5 mM beta-mercaptoethanol, and 10 mM EDTA) followed by centrifugation at 16,000 rpm for 10 minutes at 4 ºC. The inclusion bodies were then resuspended in 250 mL of denaturation buffer (20 mM Tris-HCl pH 8.5, 6 M Urea, and 100 mM beta-mercaptoethanol) and incubated at room temperature overnight. The resuspension was then centrifuged at 18,000 rpm for 10 minutes at 4 ºC and the supernatant was collected. The supernatant was diluted to about 1 mg/mL in denaturation buffer and refolded by dialysis in the following steps at 4 ºC:

1. 4 hours in 5 L of refolding buffer 1 (50 mM Tris-HCl pH 8.5, 150 mM NaCl, 5 mM CaCl_2_ , 0.5 mM ZnCl_2_ , 0.4 M sucrose, 4 M urea, 5 mM reduced glutathione, and 0.5 mM oxidized glutathione).
2. 12 hours in 5 L of refolding buffer 2 (50 mM Tris-HCl pH 8.5, 150 mM NaCl, 5 mM CaCl_2_ , 0.5 mM ZnCl_2_ , 0.4 M sucrose, 2 M urea, 5 mM reduced glutathione, and 0.5 mM oxidized glutathione).
3. 4 hours in 5 L of refolding buffer 3 (50 mM Tris-HCl pH 8.5, 150 mM NaCl, 5 mM CaCl_2_ , 0.5 mM ZnCl_2_ , 0.4 M sucrose, 5 mM reduced glutathione, and 0.5 mM oxidized glutathione)

After the final refolding step, the protein was centrifuged at 16,000 rpm for 10 minutes at 4 ºC and the supernatant was collected for purification. The protein was concentrated and injected into a HiLoad 26/600 Superdex 75 column run in 20 mM Tris-HCl pH 7.5, 5 mM CaCl_2_ , 0.5 mM ZnCl_2_ , and 150 mM NaCl at a flow rate of 2 mL/min. Peak fractions corresponding to monomeric MMP12 were collected, pooled, concentrated to <1 mg/mL, snap frozen in liquid nitrogen, and stored at -80 ºC until used.

#### Protein Crystallization

Crystals of MMP12 (G106-G263) bound to both peptide **13** and peptide **51** were obtained by incubating 0.55 mg/mL MMP12 with 0.1 mM of either peptide for 1 hour on ice. After incubation, the MMP12/peptide solution was concentrated to 8 mg/mL with a final DMSO concentration of less than 1%. For crystallization of MMP12 bound to peptide **13**, 1 µL of 8 mg/mL MMP12 (G106-G263)/peptide **13** complex was combined with 1 µL of 0.2 M lithium sulfate, 0.1 M Tris-HCl pH 8.5, 1.26 M ammonium sulfate and incubated over 500 µL of the same crystallization well solution in a hanging drop format at 18 °C. Crystals were cryo-protected in well solution supplemented with 30% glycerol, mounted on loops, and flash frozen in liquid nitrogen. Crystals of MMP12 (G106-G263) bound to peptide **51** were obtained by combining 1 µL of 8 mg/mL MMP12 (G106-G263)/peptide **51** complex with 1 µL of 0.1M Bis Tris propane pH 7.3, 1.5 M lithium sulfate. Crystals were grown in hanging drops at 18 °C. Crystals were cryo-protected in 25% ethylene glycol and flash-frozen in liquid nitrogen.

#### Crystallography data collection, processing, and structure solution

Data was collected at beamline P13 at the European Molecular Biology Laboratory synchrotron source. Data collection and processing was completed by Helix Biostructures. The data set for MMP12 bound to peptide **13** was indexed and integrated with XDS^68^ into P1 space group and scaled with Aimless^69^ . The structure was solved using PDB 1jiz^41^ as a search model with Phaser^70^ with 12 copies of the protein in the asymmetric unit. After an initial round of refinement in phenix.refine^71^ clear electron density was present for the bound peptide. Restraints for threonine hydroxamate were generated using eLBOW^72^ and the rest of peptide **13** was built using canonical amino acids in Coot^73^ followed by another round of phenix.refine. The model then underwent multiple cycles of manual building adjustment followed by automated refinement. The structure included numerous sulfate ions that were obtained from the lithium sulfate-based condition. Waters were included in the structure after confidently fitting all protein, ligand, and solvent molecules.

The data set for MMP12 bound to peptide **51** was indexed, integrated, and scaled with DIALS^74^ into space group P4_1_312. The structure was solved with Phaser using PDB 1jk3^75^ as a search model with 1 copy of the protein in the asymmetric unit. The structure was otherwise completed in a similar manner as described for the peptide **13** bound MMP12 structure.

#### Molecular Dynamics Simulations

The MMP12–peptide complex was parameterized using the AMBER ff14SB force field for the protein and peptide components^76^ . Parameters for the nonstandard hydroxamate moiety were generated using the General AMBER Force Field 2 (GAFF2), with atomic partial charges assigned using the AM1-BCC charge model in AmberTools22^77,78^ . The complex was solvated in an orthorhombic periodic box with initial dimensions of 62.0 × 65 × 60 Å^3^ using explicit TIP3P water^79^ . The final system contained 7,045 water molecules, 22 Na^+^ ions, and 23 Cl^-^ ions, corresponding to 150 mM monovalent salt while maintaining overall charge neutrality.

Two Zn^2+^ ions were retained in their experimentally defined coordination environments. To preserve metal coordination during the simulations, each Zn^2+^ ion was harmonically restrained to the three surrounding coordinating histidine residues using six Zn–His pair restraints with an equilibrium distance of 2.0 Å and a force constant of 5,000 kJ mol^-1^ nm^-2^ .

The solvated system was first subjected to steepest-descent energy minimization for up to 5,000 steps, with a maximum-force convergence criterion of 1,000 kJ mol^-1^ nm^-1^ . The system was subsequently equilibrated through six sequential restrained MD stages in which structural restraints were progressively relaxed over 1 ns.

Following equilibration, all staged positional restraints were removed and a 1-μs production MD simulation was performed in the NPT ensemble, while the Zn–His coordination restraints described above were retained. Temperature was maintained at 310 K using the velocity-rescaling thermostat with a coupling constant of 1.0 ps^80^ , and pressure was maintained isotropically at 1.0 bar using the Parrinello–Rahman barostat^81^ with a coupling constant of 5.0 ps and compressibility of 4.5 × 10^-5^ bar^-1^ .

Short-range electrostatic and van der Waals interactions were evaluated using a 10-Å cutoff. Long-range electrostatic interactions were calculated using the particle-mesh Ewald (PME) method^82^ . Van der Waals interactions employed the Verlet cutoff scheme with the potential-shift modifier, and analytical long-range dispersion corrections were applied to the energy and pressure. Bonds involving hydrogen atoms were constrained using the LINCS algorithm^83^ , permitting an integration time step of 2 fs. All equilibration and production simulations were performed using GROMACS 2023^84^ .

#### Structure analysis and visualization

Protein structure figures were generated with PyMol software version 3.1.3 (Schrödinger, LLC).

## Contributors

Vladimir Akhmetov, Emily Kustra, Amirhossein Mafi, James Mulligan, Isaac Knudson, Noah Durham, Manjot Sangha, Bryan Argueta†, Stephanie Fullam, Zhenghao Chen, Janani Sridar, Michelle Tran, John C.K. Wang, Qi Hao, Jun Xu, Jeffmin Lin, and Matthew Bratkowski †Deceased

## Acknowledgments

We thank Houxia Shi and the team at Biortus for help with the expression and purification of MMP12. We thank Josh Carter, Ann Mulichak, Thomas Sawyer and team members at Helix Biostructures for data collection and processing of MMP12 structures. We thank WuXi Apptec for peptide pharmacokinetic studies. The synchrotron data was collected at the beamline operated by EMBL Hamburg at the PETRA III storage ring (DESY, Hamburg, Germany). We would like to thank Saravanan Panneerselvam and staff for assistance in using the beamline. We thank Chuan Sheng Foo, Meghana Kshirsagar, Dan Eaton, Snahel Patel, and Matt Onsum from Calico Life Sciences LLC for reviewing the manuscript and providing helpful suggestions.

## Competing Interests

This work was funded by Calico Life Sciences LLC. All contributors are current or former employees or contractors at Calico Life Sciences LLC.

