## Supplemental Information for "Accelerated discovery and optimization of a potent and selective MMP12 inhibitor with the machine learning platform CAMOLE-1"

Calico Life Sciences LLC

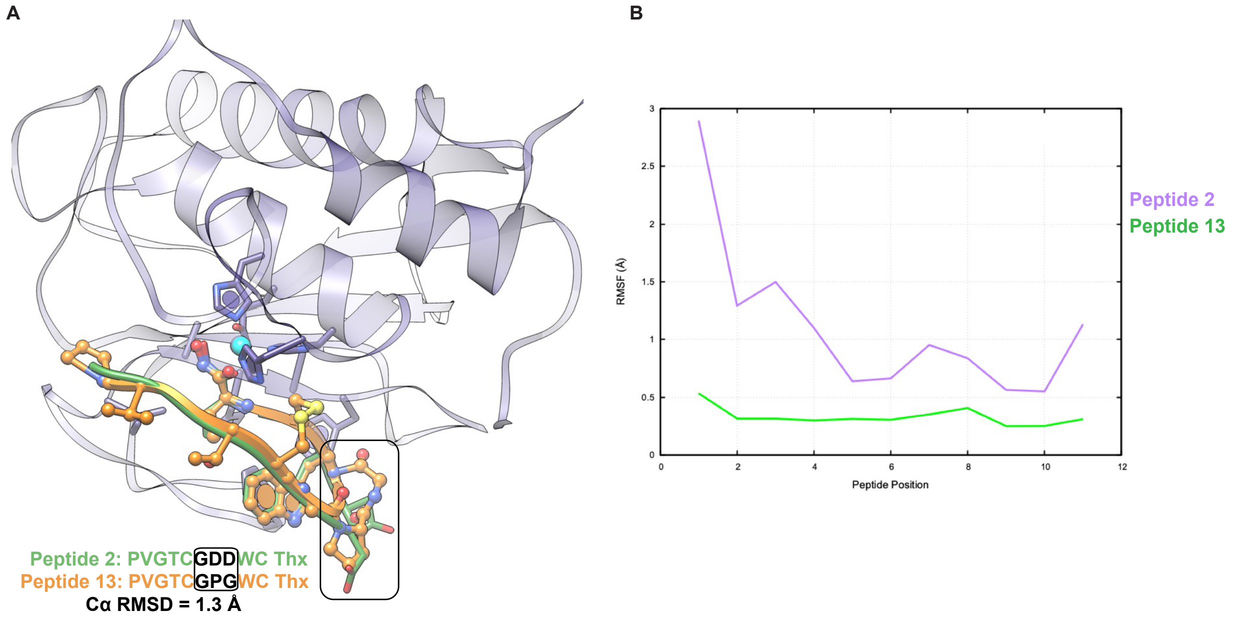

**Figure S1: MD simulations used to optimize initial peptide potency**

A) Computational models of peptides **2** and **13**. B) 1 μs MD simulations indicated backbone fluctuations for peptide **2**. In contrast, peptide **13** was much more stable.

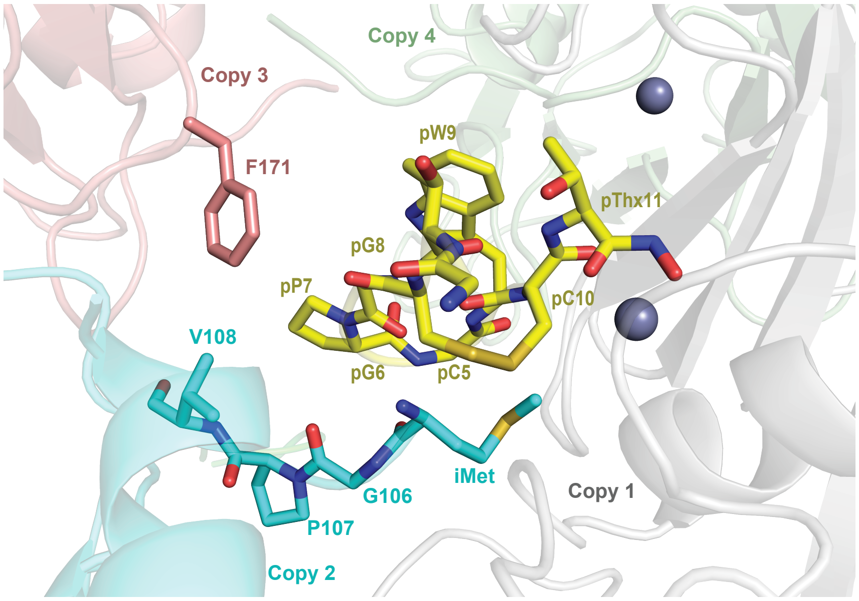

**Figure S2: Crystal packing interactions for the MMP12 + peptide 13 structure**

Packing interactions between different copies of MMP12 in the asymmetric unit (*cyan, salmon, and green cartoon*) in relation to peptide **13** (*yellow sticks*) bound to MMP12 copy 1 (*gray cartoon*). MMP12 residues close to peptide **13** in crystal packing interactions are show as sticks. Here "iMet" refers to the non-native initiator methionine at the N-terminus of the protein.

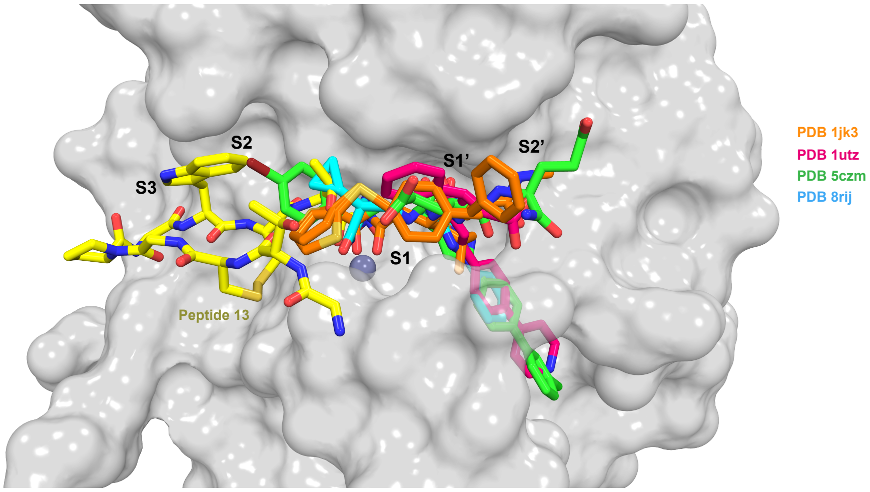

**Figure S3: Structural comparison of peptide 13 binding with other inhibitors**

Structures of MMP12 bound to the hydroxamate inhibitor batimastat (PDB 1jk3[1], *orange sticks*), bound to the non-zinc chelator PF-00356231 (PDB 1utz[2], *raspberry sticks*), bound to the peptide-like compound RXP470 (PDB 5czm[3], *green sticks*), and bound to the hydantoin chelator compound 3 (PDB 8rij[4], *cyan sticks*) were aligned to our structure bound to peptide **13** (*yellow sticks*). MMP12 is shown as a semi-transparent surface to best illustrate the S and S′ subsites.

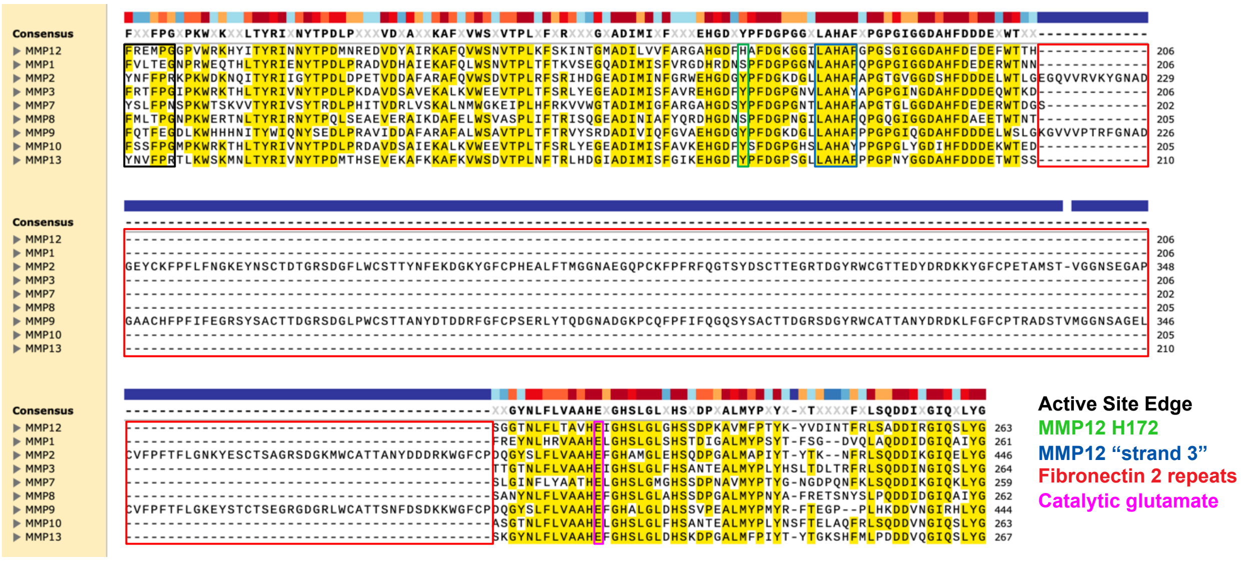

**Figure S4: Sequence alignment of the active site of different MMPs**

UniProt[5] sequences for the human catalytic domains of MMP12 (P39900), MMP1 (P03956), MMP2 (P08253), MMP3 (P08254), MMP7 (P09237), MMP8 (P22894), MMP9 (P14780), and MMP10 (P09238) were aligned using a Clustal Omega[6] based alignment in SnapGene software (Dotmatics, www.snapgene.com). Regions discussed in the report are boxed and identified by color-coding.

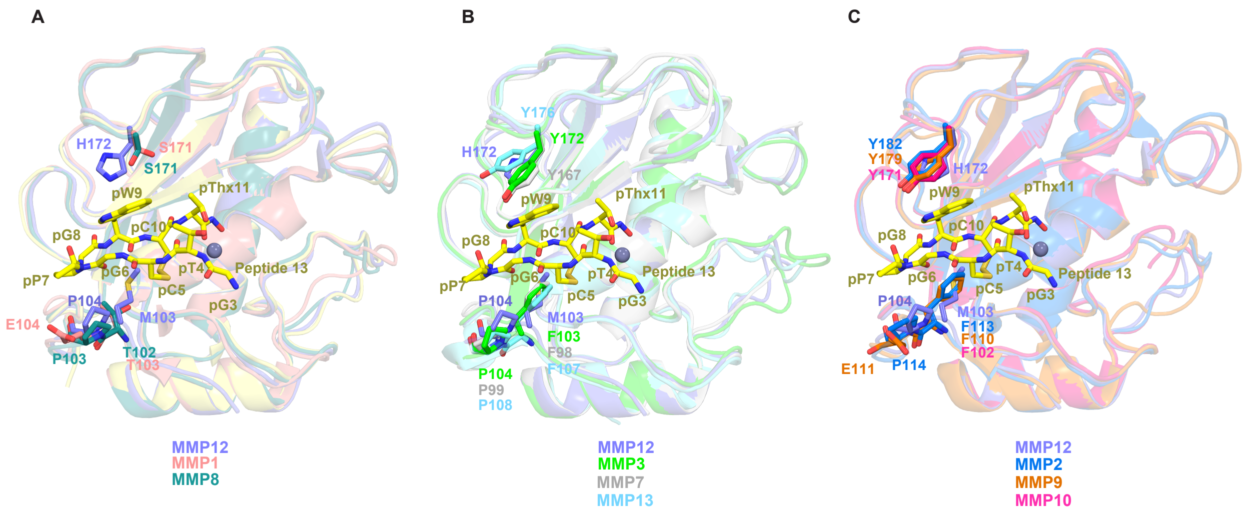

**Figure S5: Alignment comparison of other MMPs with the peptide 13-bound MMP12 structure**

Our MMP12/peptide **13** crystal structure (peptide in yellow sticks) was first aligned to a slightly longer structure of MMP12 (PDB 1jiz[7], *purple cartoon*) for the purpose of hypothesizing the molecular mechanism of selectivity observed from activity data. Published structures of the catalytic domains of other MMPs were then compared by alignment to the MMP12/peptide **13** structure. For simplicity, bound ligands for other MMP structures are not shown. A) Alignment with structures of MMP1 bound to a triple helical collagen peptide (PDB 4auo[8], *salmon*) and MMP8 bound to a peptidomimetic hydroxamate compound (PDB 1kbc[9], *teal*). B) Alignment with structures of MMP3 bound to an N-carboxy-alkyl inhibitor (PDB 1sln[10], *green*), MMP7 bound to a peptide-small molecule hybrid (PDB 8jug[11], *gray*), and MMP13 bound to an L-glutamate motif inhibitor (PDB 3tvc[12], *cyan*). C) Alignment with structures of MMP2 bound to a decapeptide inhibitor (PDB 3ayu[13], *blue* ), MMP9 bound to a hydroxamate-based inhibitor (PDB 4h3x[14], *orange*), and MMP10 bound to hydroxamate inhibitor (PDB 1q3a[15], *pink*).

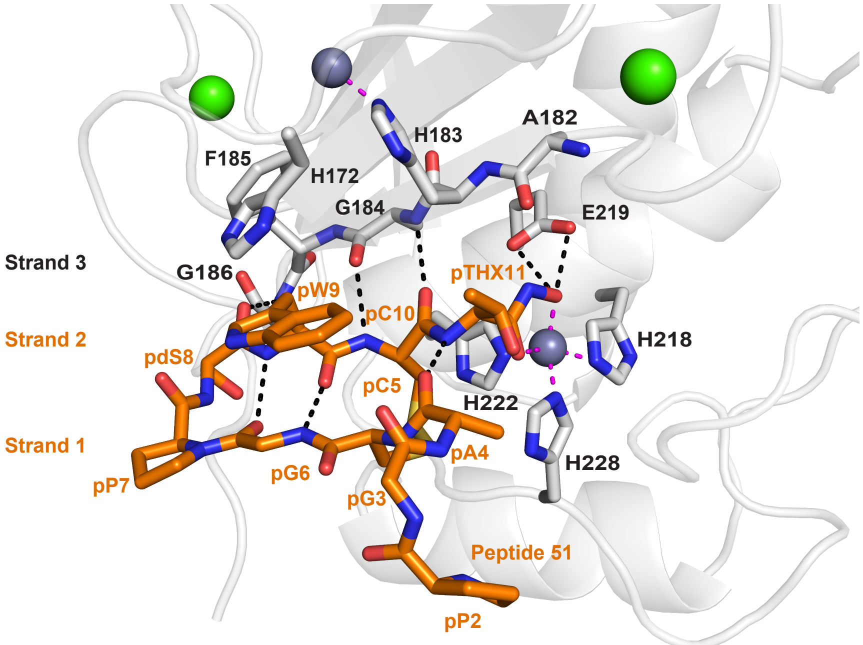

**Figure S6: Detailed interactions between MMP12 and peptide 51 from the crystal structure**

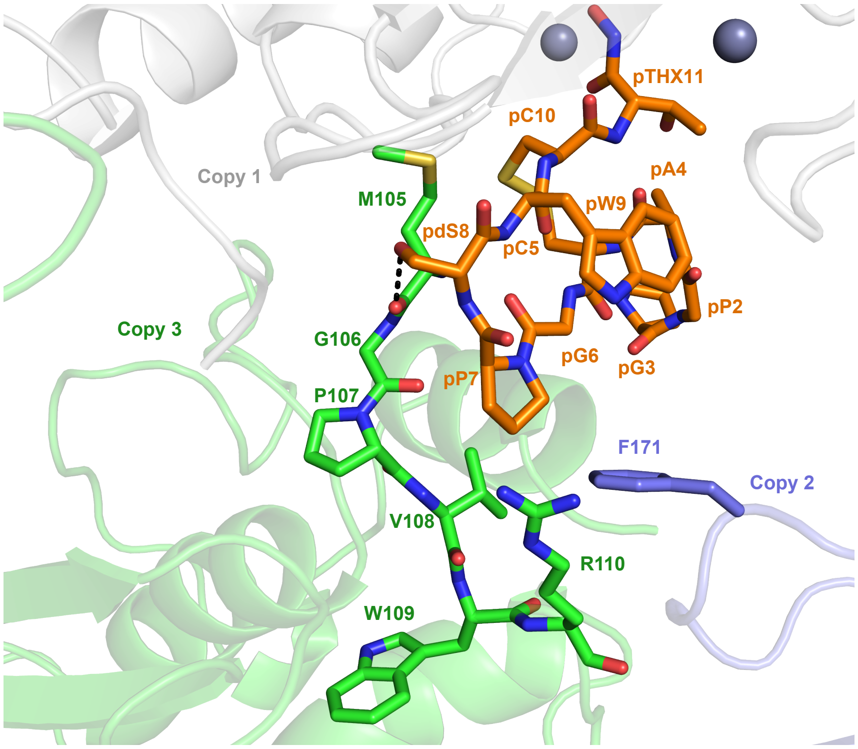

**Figure S7: Crystal packing interactions for the MMP12/peptide 51 structure**

Packing interactions between different copies of MMP12 in the asymmetric unit (*purple and green cartoon*) in relation to peptide **51** (*orange sticks*) bound to MMP12 copy 1 (*gray cartoon*). MMP12 residues close to peptide **51** in crystal packing interactions are shown as sticks. While peptide **51** interacts with additional residues in symmetry mates, these are likely crystal packing artifacts without physiological relevance.

| | Peptide 13 | Peptide 51 |
|---|---|---|
| **Data collection** | | |
| Space group | P1 | $P4_132$ |
| Cell dimensions | | |
| $a$, $b$, $c$ (Å) | 81.94, 84.75, 93.35 | 108.63, 108.63, 108.63 |
| $\alpha$, $\beta$, $\gamma$ (°) | 73.24, 67.24, 70.78 | 90.00, 90.00, 90.00 |
| Resolution (Å) | 44.64 – 1.86 (1.91 – 1.86)* | 44.35 – 2.59 (2.64 – 2.59) |
| $R_{merge}$ | 0.106 (0.692) | 0.132 (1.31) |
| $I / \sigma I$ | 7.6 (1.7) | 14.3 (1.1) |
| $CC_{1/2}$ | 0.994 (0.351) | 0.996 (0.836) |
| Completeness (%) | 94.2 (93.6) | 100 (100) |
| Redundancy | 3.7 (3.8) | 22.5 (24.3) |
| **Refinement** | | |
| Resolution (Å) | 44.64 – 1.86 | 44.35 – 2.59 |
| No. reflections | 170095 | 7259 |
| $R_{work}$ / $R_{free}$ | 0.1897 / 0.2324 | 0.244 / 0.290 |
| No. non-hydrogen atoms | | |
| Protein and peptide | 15592 | 1316 |
| Non-amino acid ligand† | 346 | 35 |
| Solvent | 2096 | 0 |
| $B$-factors (Å$^2$) | | |
| Protein and peptide | 25.49 | 82.29 |
| Non-amino acid ligand† | 32.15 | 88.49 |
| Solvent | 33.86 | 0 |
| R.m.s. deviations | | |
| Bond lengths (Å) | 0.005 | 0.004 |
| Bond angles (°) | 0.667 | 0.720 |
| Ramachandran plot† | | |
| Favored (%) | 97.69 | 98.14 |
| Allowed (%) | 2.31 | 1.86 |
| Outliers (%) | 0.00 | 0.00 |

* Values in parentheses are for the highest-resolution shell

†Threonine hydroxamate and D-serine are treated as ligands for the purposes of these calculations. D-serine is an outlier in standard L-amino acid Ramachandran plots.

**Table S1: Data collection and refinement statistics**

| Peptide | Sequence | MMP12 IC50 (nM) |
|---|---|---|
| **17** | ATGACGPGWC[Thx] | 4.94 |
| **18** | NTGACGP[dP]WC[Thx] | 7.3 |
| **19** | S[Cpa]GACGP[dS]WC[Thx] | 6.55 |
| **20** | ATGACGS[dP]WC[Thx] | 2.94 |
| **21** | GPGACG[NMeI][dP]WC[Thx] | 10.6 |
| **22** | APG[dS]CGP[dI]WC[Thx] | 9.95 |
| **23** | S[Hyp]GACGP[dP]WC[Thx] | 5.02 |
| **24** | [dL]PGACGA[dP]WC[Thx] | 5.01 |
| **25** | [dL]PGGCG[NMeA][dP]WC[Thx] | 2.29 |
| **26** | N[Hyp]GGCGP[dP]WC[Thx] | 3.75 |
| **27** | SP[dS]ACGP[dP]WC[Thx] | 1.46 |
| **28** | [NMeD]PGACGA[dP]WC[Thx] | 3.79 |
| **29** | [NMeA]TGACGP[dP]WC[Thx] | 5.07 |
| **30** | TAG[dS]CGPGWC[Thx] | 1.82 |
| **31** | SPGGCGT[dH]WC[Thx] | 5.19 |
| **32** | DVGGCGP[dP]WC[Thx] | 1.01 |
| **33** | SVGACGP[dP]WC[Thx] | 1.19 |
| **34** | [dV]TGACGP[dP]WC[Thx] | 8.18 |
| **35** | [Cpa][Thz]GACGP[dP]WC[Thx] | 4.76 |
| **36** | [Cpa][Hyp]GGCGP[dP]WC[Thx] | 5.59 |
| **37** | SPGACGPGWC[Thx] | 4.67 |
| **38** | SIGACGP[dP]WC[Thx] | 2.52 |
| **39** | [Cval][Hyp]GGCGP[dP]WC[Thx] | 2.84 |
| **40** | S[Nip]GACGP[dS]WC[Thx] | 4.5 |
| **41** | EPGACGP[dP]WC[Thx] | 3.4 |
| **42** | SPGTCGP[dP]WC[Thx] | 11.2 |
| **43** | A[Thz]GACGT[dP]WC[Thx] | 9.95 |
| **44** | A[NMeA]GACGP[dP]WC[Thx] | 7.17 |
| **45** | ATGACG[Hyp][dP]WC[Thx] | 8.6 |
| **46** | S[NMeT]GACGP[dP]WC[Thx] | 4.93 |
| **47** | NPGACGA[dP]WC[Thx] | 7.17 |
| **48** | [Cval]TGACGP[dP]WC[Thx] | 6.92 |
| **49** | SPGACGP[dP]WC[Thx] | 7.7 |
| **50** | A[Hyp]GGCG[NMeT]GWC[Thx] | 2.82 |
| **51** | [dS]PGACGP[dS]WC[Thx] | 2.21 |
| **52** | APGGCG[NMeA][dP]WC[Thx] | 2.92 |
| **53** | [dS]PGGCG[NMeA][dP]WC[Thx] | 5.2 |
| **54** | [NMeDAla][Thz]GACGP[dP]WC[Thx] | 6.48 |
| **55** | STGACGP[dP]WC[Thx] | 5.07 |

**Table S2: MMP12 IC50 potencies for ML-optimized peptides**

Abbreviations are as follows: "d" (D-amino acid), "NMe" (N-methylated amino acid), "Cpa" (cyclopropyl-alanine), "Hyp" (trans-L-hydroxy-proline (2S, 4R)), "Thz" (thiazolidine carboxylic acid), "Cval" (cyclovaline AC4C), "Nip" (nipecotic acid), and "NMeDAla" (N-methylated D-Alanine).
